# Draft genome analysis of *Delftia tsuruhatensis* IICT-RSP4, a strain with uricase potential isolated from soil

**DOI:** 10.1101/2024.05.08.592295

**Authors:** Mahesh Anumalla, Uma Rajeswari Batchu, Bhima Bhukya, Prakasham Reddy Shetty

## Abstract

*Delftia tsuruhatensis* IICT-RSP4, an uricase producing bacterium was isolated using *i*-chip method from soil and characterized. Here, we report the draft genome sequence of *D. tsuruhatensis* IICT-RSP4. The genome data comprised of 6,627,718bp (6.6 MB) with a GC content of 66.6% with 7 protein encoding genes, 346 sub-systems with 6165 coding sequences and 112 RNAs. The genome revealed five functional secondary metabolite biosynthetic gene clusters viz. terpene, resorcinol, NRP+PKS, T2PKS, and RiPPS related to antimicrobial, anticancer and antimalarial functionality. *In vitro*, screening studies revealed the uricase potential of the strain *D. tsuruhatensis* IICT-RSP4. This is the first report on the whole genome sequence of an uricase-producing *D. tsuruhatensis* IICT-RSP4.

## Introduction

*Delftia tsuruhatensis* is a ubiquitous bacterium with high genetic diversity and inhabit various ecological niches (Yin et al. 2022). It was previously isolated from rhizosphere soil, activated sludge, and polluted environments. In Russia, *D. tsuruhatensis* MR-6/3H strain with multi-drug resistant potential was reported from raw bovine milk (Andriyanov et al. 2024). It is a gram-negative bacterium with versatile functions wherein to promote plant growth in rhizosphere soil (Han et al. 2005), degradation oforganic pollutants (Zhang et al. 2010) and is also considered as an opportunistic human pathogen (Preiswerk et al. 2011). In addition, it is also able to produce substances to inhibit plant and human pathogens (Hou et al. 2015; Tejman et al. 2019). In the present study, strain *D. tsuruhatensis* IICT-RSP4 was isolated from a soil sample collected from the Jalavihar area of Indian Institute of Chemical Technology, India. The primary *in vitro* screening studies revealed the uricase potential of the strain using uric acid agar plate assay by measuring the zone of clearance due to degradation of uric acid around the colony (Lehejckova et al. 1986). Uricase is an essential enzyme that catalyzes the degradation of uric acid into water soluble allantoin to prevent toxic accumulation of uric acid, thus used in the treatment of refractory gout and tumor lysis syndrome (George and Sundy 2012). Hence, the present study explored the draft genome sequence analysis of *D. tsuruhatensis* IICT-RSP4, a new strain with uricase potential. The analysis was performed using next-generation sequencing and phylogenomic approaches. So far, this is the first report on the draft genome sequence of uricase positive *D. tsuruhatensis* IICT-RSP4.

## Results

### Isolation and identification of bacteria

Soil samples were collected in self-sealing covers from different locations of the Jalavihar area of CSIR-Indian Institute of Chemical Technology (CSIR-IICT), Hyderabad, India and preserved in cold cabinet (4 ºC) until further use. The collected soil samples were screened for novel microbes using previously adopted *i*-Chip method in the lab (Tejaswi et al. 2023).

### Screening of isolates for uricase production

A total of 4 *i*-Chip isolates were screened for uricase production by using the basal uricase screening media consisting of uric Acid-5g/L, glycerol-30g/L, NaCl-5g/L, K_2_HPO_4_-1g/L, MgSO_4_. 7 H_2_O-0.2g/L and CaCl_2_-0.1g/L. Briefly, 100 μl cell free extract of the selected isolates was added into 6mm wells punched in the uric acid screening agar plate and incubated at 30 ºC for 24-48 hrs to observe the zone of uric acid clearance using well diffusion assay. The potential uricase producing strain was selected based on the diameter of the zone of clearance. (Abdel-Fattah & Abo-Hamed 2002). The results revealed that among all four isolates uricolytic potential was detected by bacterial isolate IICT-RSP4 based on the development of clear zone around the colony on uric acid agar plate (Fig1).

**Fig1.**
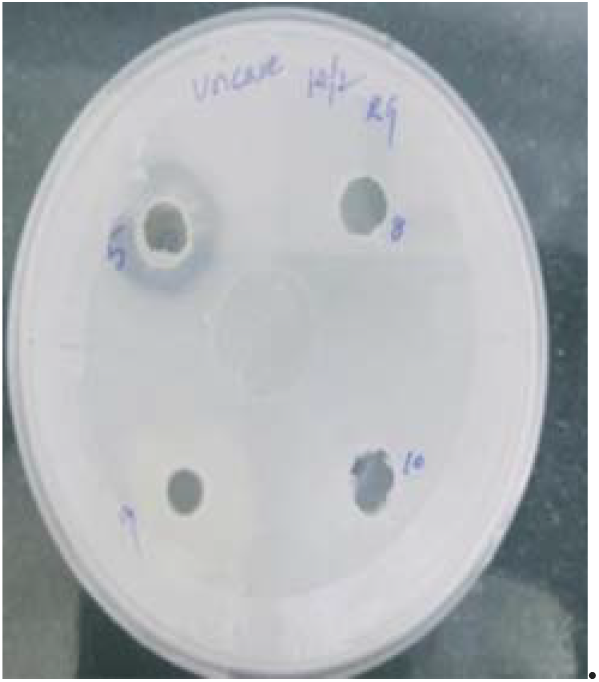
Uricase potential of IICT-RSP4 showing clear zone around the culture supernatant of selected isolate in agar well diffusion assay.

### DNA isolation, sequencing and functional genome analysis

DNA was extracted from the pure culture using a HiPurA Bacterial Genomic DNA Purification kit (MB505-250PR HiMedia Laboratories PVT Ltd, Nashik, India) according to the manufacturer’s instructions. The whole genome sequence was characterized by Nano pore sequencing technology on a flow cell FLO-MIN114 and kit type SQK-LSK114 using guppy base caller (version 6.4.6). The base calling was done using the super accuracy model DNAr (10.4.1_e82_400bps_sup.cfg). The resulting fastq file was quality trimmed using default parameters of Porechop (version 0.2.4) (Wick et al. 2017) to remove low quality sequences (<2 kb) and assembled into contigs with Flye version 2.9 (Kolmogorov et al. 2020). The trimmed contig data was used for taxonomic classification using kraken 2 version 2.0.7 (Wood et al. 2019).

The analysis of the primary draft genome was performed by rapid annotation using subsystems technology (RAST) (Brettin et al. 2015). The functional genome analysis using RAST explored that the genome consists of 7 contigs with a total contig size of 6,627,718bp (6.6 MB). The N50 contig number was1044867 and L50 value was 2. The G+C content of the genome was 66.6 % with 7 protein encoding genes, 346 sub-systems with 6165 coding sequences (Table 1, Fig2), and 112 RNAs.

**Fig2.**
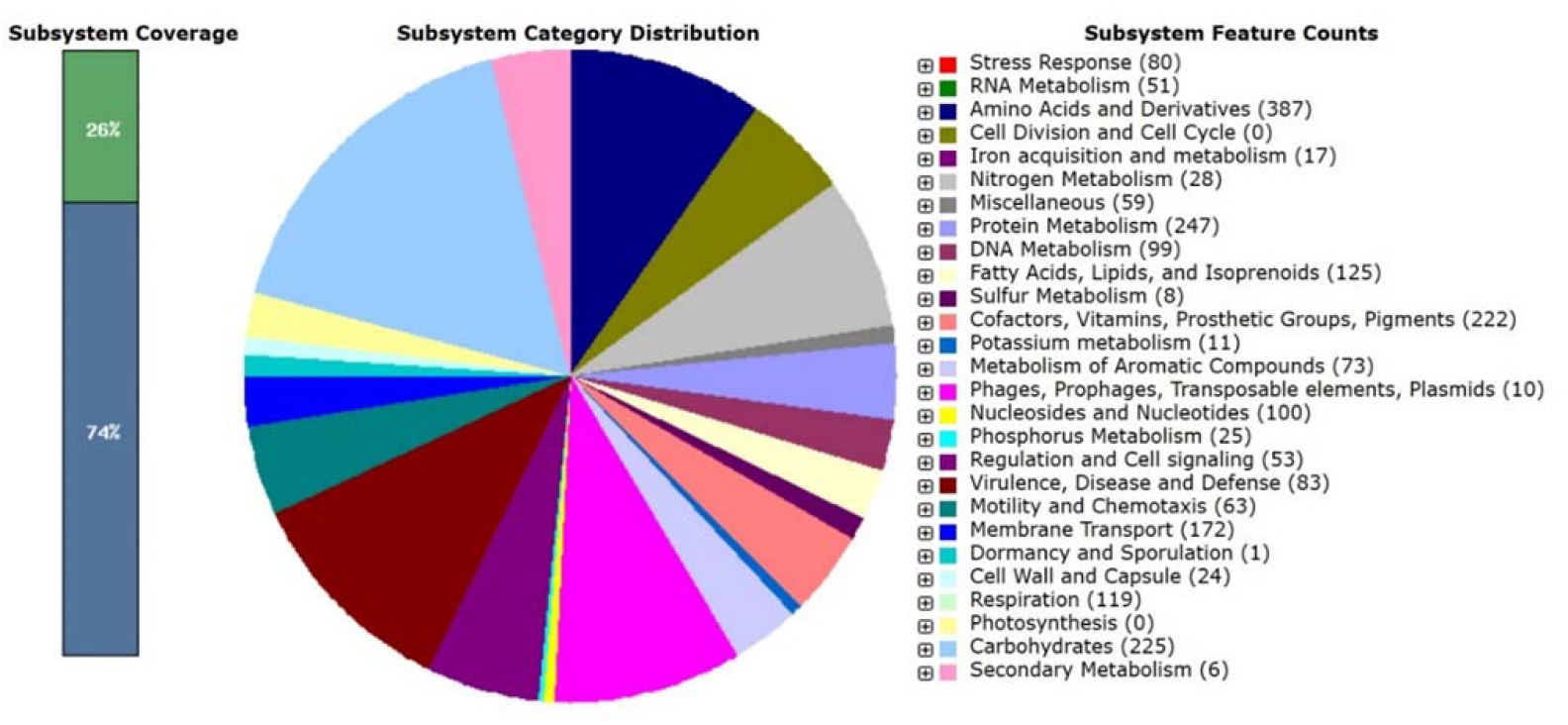
Sub-system statistics information of *D. tsuruhatensis* IICT-RSP4 using RAST annotation. The category of subsystems and corresponding feature counts were displayed in the legend.

**Table 1:** Draft genome characteristics of *D. tsuruhatensis* IICT-RSP4.

|  |  |
| --- | --- |
| Number of contigs | 7 |
| Size of contig (bp) | 6,627,718 |
| N50 contig number | 1044867 |
| L50 | 2 |
| G+C content (%) | 66.6 |
| Contigs with protein-encoding genes | 7 |
| No. of subsystems | 346 |
| No. of coding sequences | 6165 |
| No. of RNAs | 112 |

Further, the *D. tsuruhatensis* IICT-RSP4 draft genome was analyzed for secondary metabolite biosynthetic gene clusters using antiSMASH version 5.0 (Blin et al. 2019). The analysis identified five secondary metabolite biosynthesis gene clusters i.e. terpene, resorcinol, NRP+PKS, T2PKS, and RiPPS (Table 2). Among these, terpene, resorcinol, and NRP+PKS are frequently identified genes in *D. tsuruhatensis*, whereas T2PKS and RiPPS are rarely identified genes that can be obtained by horizontal transfer mechanisms from other species. Terpenes can exhibit anticancer and antimalarial potential (Kamran et al. 2022: Gabriel et al. 2018). The antiSMASH homology data represents that NRP+PKS and T2PKS contained homologous genes with known clusters. The non-ribosomal peptide (NRP)-metallophore (NRP+PKS) with delftibactin A (A biosynthetic gene cluster from *D. acidovorans* SPH-1) showed 100% homology. Delfibactins are secondary metabolites belonging to the group of siderophores which can detoxify the soluble gold ions (Sussmuth and Mainz 2017). NRP-Delfibactins are demonstrated as an antimicrobial agent against multi-drug resistant bacteria like MRSA and VRE as well as *Acinetobacter baumannii* (Tejman et al. 2019). Whereas, the type-II polyketide synthase (T2PKS) biosynthetic gene cluster shows only 10% homology with pyxidicycline (A biosynthetic gene cluster from *Pyxidicoccus fallax*). Pyxidicyclines are natural products that belong to a class of topoisomerase inhibitors to target replication of oncogenes and pathogens (Panter et al. 2018). It might represent the finding of novel metabolite genes with potential biotechnological applications like the promotion of plant growth and bio-controlling ability.

**Table 2:** antiSMASH annotation of the draft genome of *D. tsuruhatensis* IICT-RSP4 for identification of secondary metabolites.

| Features | Number of clusters |
| --- | --- |
| No of smCOG<sup>1</sup> | 5 |
| PKS<sup>2</sup> |  |
| T2PKS (Type II) | 1 |
| NRPS<sup>3</sup> |  |
| NRPS- T1PKS (Type I) | 1 |
| Biosynthetic Pathways |  |
| Resorcinol | 1 |
| Terpenes | 1 |
| RiPP-like | 1 |

Note: 1. Secondary metabolism Clusters of Orthologous Groups 2.Polyketide synthase 3.Nonribosomal polypeptide synthetase.

However, no homology was observed for other gene clusters in the antiSMASH data base, especially for ribosomally synthesized and post-transitionally modified peptides (RiPPS) which are biologically potent compounds of natural origin (Ozaki et al. 2023).

The Kraken2 output observed that 98% of the reads were classified as bacteria and only 2% were left unclassified. Overall, the classification based on the ‘standard-8 database’ yielded a single species of *Delftia* in the sample sequenced and was identified as a species of *D. tsuruhatensis* with a similarity of 80% (Fig3). To eliminate the potential false positives, a species richness cut-off of >1% was considered and was considered the true result.

**Fig3.**
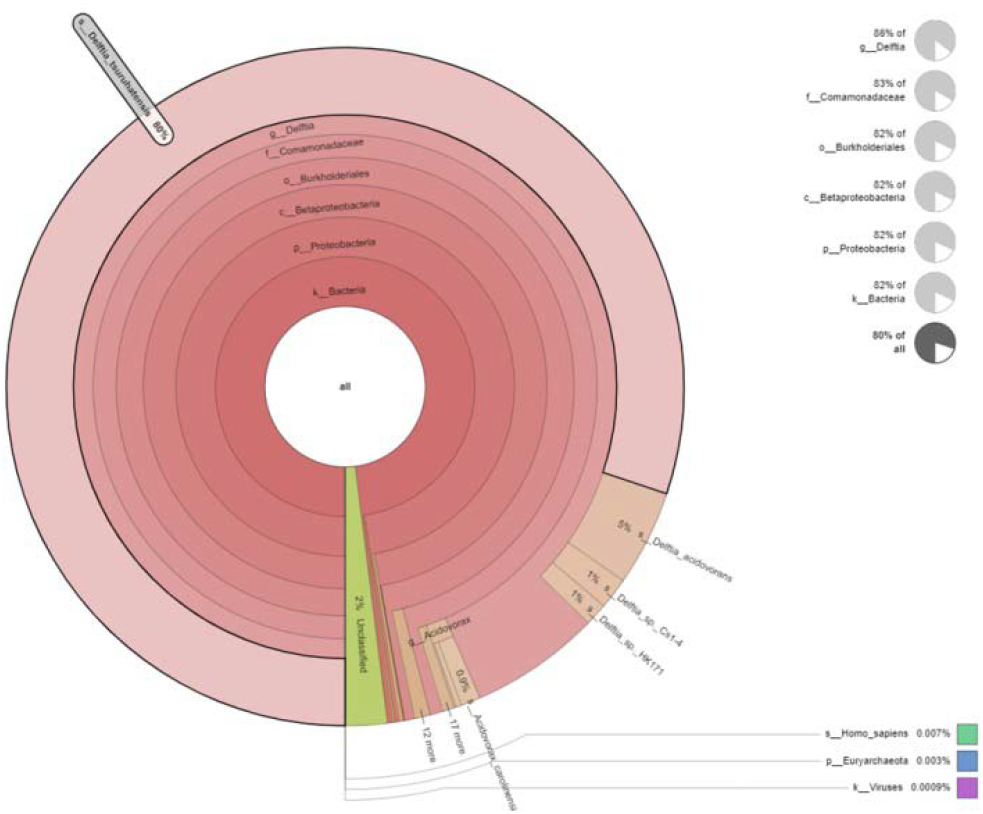
Krona graph representing the DNA sequence reads of *D. tsuruhatensis* IICT-RSP4

### 16s rRNA sequencing

BLASTN version 2.12.0 was used to carry out basic local alignment search tool (BLAST) analysis (Altschul et al. 1990) against NCBI database for species-level identification. The top 20 near species 16S rRNA gene sequence was retrieved. However, these results were failed to delineate between *Delftia* species. Hence, 16s rRNA gene comparison was performed by Type (Strain) Genome Server (TYGS) (Meier et al. 2019) and Genome BLAST Distance Phylogeny (GBDP) tree was constructed using 16s rRNA gene and whole genome sequence. In addition, Average Nucleotide Identity (ANI) between related species with whole genome was calculated by using OrthoANIu algorithm (Yoon et al. 2017).

BLASTn alignment (version 2.12.0) with the 16S rRNA sequence database showed 100% sequence similarity with *D. tsuruhatensis* BN-HKY1 (Gene accession number HQ731448.1) and 99.84% sequence similarity with *D. lacustris* (Gene accession number MH675503.1). The 16S rRNA gene sequence has been deposited in the NCBI under the Gene accession number PP3969.

The results of TYGS showed that 16s rRNA gene sequence of IICT-RSP4 strain belongs to the same clade of *D. tsuruhatensis* NBRC 16741 and *D. lacustris* LMG 24775 (Fig4). The % dDDH threshold values > 70% and diff G+C content <1% from TYGS analysisandANI value > 95-96% indicates that the strain IICT RSP4 belongs to *D. tsuruhatensis* (Table 3). Though, different species *D. tsuruhatensis* and *D. lacustris* belongs to single clade (clade A) whereas *D. acidovorans* belongs to different clade (clade C).

**Fig4.**
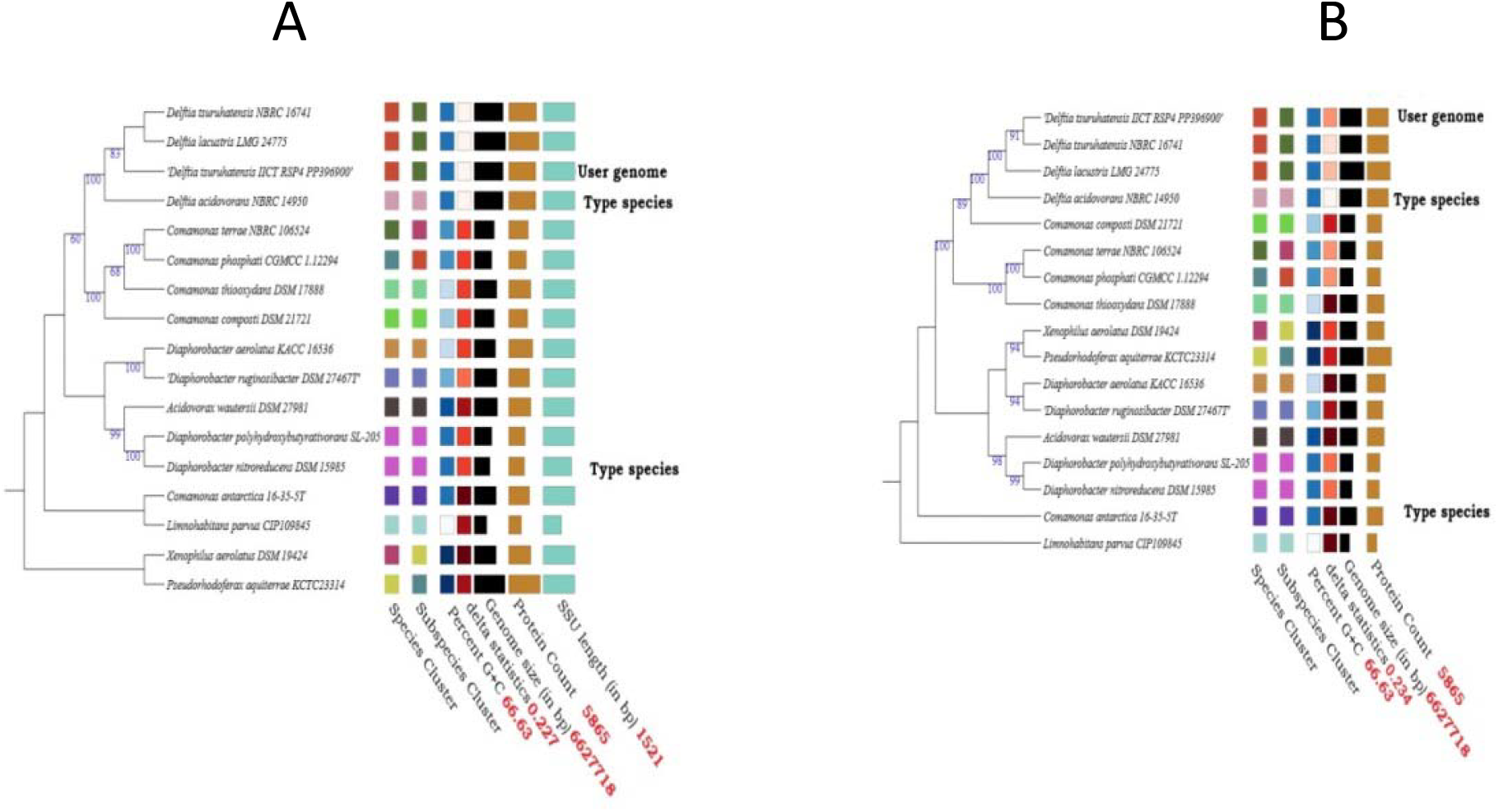
Genome BLAST Distance Phylogeny method (GBDP) diagram of *D. tsuruhatensis* IICT-RSP4 generated using TYGS shows that *D. tsuruhatensis* IICT RSP4 was closely related with the *D. tsuruhatensis* A) Phylogram based on 16S rRNA gene sequence B) Phylogram based on Whole genome sequence

**Table 3:** Digital DNA hybridization (dDDH), ANI values and % G+C content of *D. tsuruhatensis* IICT-RSP4 with its closely related species using TYGS analysis.

| Species Name | dDDH (%) | ANI | Diff G+C% |
| --- | --- | --- | --- |
| *Delftia tsuruhatensis* NBRC 16741 | 88.0 | 98.78 | 0.04 |
| *Delftia lacustris* LMG 24775 | 85.7 | 99.06 | 0.41 |
| *Delftia acidovorans* UBA 7660 | 59.8 | 94.40 | 0.13 |
| *Diaphorobacter nitroreducens* DSM 15985 | 23.1 | 78.56 | 0.22 |
| *Comamonas antarctica* 16-35-5T | 23.0 | 78.55 | 0.33 |

## Acknowledgments

PRS and URB thank Council for Scientific and Industrial Research (CSIR), New Delhi for funding under the Emeritus Scheme, grant number 21(1102)/20/EMR-II and CSIR-RA fellowship, respectively. Authors thanks to Dr. Divya Tej Sowpati for his constant support for NGS sequencing. The communication number issued by CSIR-IICT for this article is IICT/Pubs./2024/139.

## Data availability

The whole genome shotgun project was deposited at DDBJ/ENA/GenBank under the accession number JBBBVL000000000 (Version JBBBVL010000000) and the Bio Project accession number was PRJNA1083484.

## Statement and Declarations

### Data Archiving Statement

The whole genome sequence has been deposited in the NCBI database under the accession number JBBBVL000000000 whereas, the 16s rRNA gene sequence was submitted under the gene accession number PP3969. The Research data will be shared after publication upon request.

